# Prolonged Hyperactivity Elicits Massive and Persistent Chloride Ion Redistribution in Subsets of Cultured Hippocampal Dentate Granule Cells

**DOI:** 10.1101/2024.10.16.618704

**Authors:** Hajime Takano, Fu-Chun Hsu, Srdjan Joksimovic, Douglas A. Coulter

## Abstract

Chloride ions play a critical role in neuronal inhibition through the activity of chloride-permeable GABA_A_ receptor channels. Ion transporters, chloride channels, and immobile ion species tightly regulate intracellular chloride concentrations. Several studies related to epilepsy suggest that chloride extrusion function may decrease in an activity-dependent manner. Consequently, it is crucial to investigate whether intense neuronal activity, as observed during status epilepticus, could lead to sustained increases in intracellular chloride levels in neurons, which in turn could contribute to epilepsy-associated hyperexcitability. This study utilized the chloride sensitive indicator (6-Methoxyquinolinio) acetic acid ethyl ester bromide (MQAE) combined with fluorescence lifetime imaging (FLIM) to examine whether application of the convulsant, pilocarpine, a muscarinic acetylcholine receptor agonist, could induce synchronous epileptiform activity and elevate intracellular chloride concentrations in hippocampal slice cultures. Using a Gaussian mixture model, we identified a multimodal distribution of intracellular chloride levels among neurons, with a significant subset of these cells exhibiting massive and prolonged (days) chloride accumulation. The combination of multicellular imaging and statistical analysis served as a powerful tool for studying the emergence of multiple, distinct populations of neurons in pathological conditions, in contrast to homogeneous populations evident under control conditions.

**Highlights:**

- Maintaining low [Cl^−^]_in_ is important for inhibitory function, however, hyperactivity, such as that seen in epilepsy, may lead to elevated [Cl^−^]_in._
- Pilocarpine induces hyperactivity in dentate granule cells (DGCs) in hippocampal organotypic slice cultures.
- Multicellular imaging using a chloride sensing dye with a fluorescence lifetime imaging approach revealed that [Cl^−^]_in_ is elevated in a subpopulation of DGCs.
- Gaussian mixture model analysis is a powerful tool for studying the emergence of cellular heterogeneity in a pathological condition.

## INTRODUCTION

Understanding how individual cells behave during various biological processes, such as development, disease progression, and therapeutic treatment, is a fundamental aspect of biological research. Traditionally, cell biology and physiology have relied on population-level measurements, assuming homogeneity in cellular responses to environmental stimuli, with the population mean representing these responses accurately. However, significant cell-to-cell variations often exist within any population (Cembrowski, 2019), meaning the population mean may not truly reflect the behavior of individual cells. In this context, our study focuses on cases where cellular heterogeneity in response to environmental changes is critical and outlines practical strategies for analyzing and interpreting this cellular heterogeneity.

In the central nervous system, chloride ion homeostasis is crucial for inhibitory mechanisms mediated by gamma-aminobutyric acid type A (GABAᴀ) receptors, which are ligand-gated chloride ion channels (Ben-Ari, 2014). The intracellular chloride ion concentration ([Cl⁻]_in_) is typically tightly regulated by combined actions of the potassium-chloride cotransporter 2 (KCC2) (Kelley et al., 2018), sodium-potassium-chloride co-transporter 1 (NKCC1) (Dzhala et al., 2010), along with other chloride channels, as well as by the extracellular and intracellular microenvironments (Glykys et al., 2014; Liu et al., 2017; Rahmati et al., 2021). In healthy adult neurons, [Cl⁻]_in_ is maintained at a low level to facilitate chloride ion influx upon GABAᴀ receptor activation, resulting in the hyperpolarization of postsynaptic neurons, controlling excitability. However, elevated [Cl⁻]_in_ levels can lead to GABAᴀ receptor-mediated chloride ion efflux, causing neuronal depolarization favoring excitability, which is implicated in the development of epilepsy (Kahle and Staley, 2008).

In epilepsy, patients typically function normally when not experiencing seizures, suggesting that even in epileptic brains, a significant proportion of individual cells can operate correctly. This observation raises the possibility that only a subset of cells exhibits abnormalities, such as elevated [Cl⁻]_in_. When triggered, these abnormal cells may dominate network properties, leading to a hyperexcitable network. For example, an earlier study of tissues from epilepsy patients found that approximately 20% of subicular pyramidal cells showed elevated [Cl⁻]_in_, despite being morphologically indistinguishable from the rest of the population (Cohen et al., 2002). Similarly, in tissues from an animal model (Pathak et al., 2007), around half of the cells investigated displayed elevated [Cl⁻]_in_ following an epilepsy-inducing stimulus. This variability within the same neuronal population in response to specific stimuli contributes to cell-to-cell heterogeneity. Understanding and controlling cellular heterogeneity is critical, particularly for manipulating subcellular populations and treating diseases (Cembrowski, 2019; Cess and Finley, 2020; Kinnunen et al., 2021).

Recent advancements in multicellular imaging techniques, such as calcium indicator probes, have enabled researchers to investigate cellular population behaviors and heterogeneity that were once thought to be homogeneous (Ikegaya et al., 2004; Allene et al., 2008; Bonifazi et al., 2009; Takano et al., 2012; Bui et al., 2015). Similar to calcium imaging, chloride imaging can assess chloride levels in many neurons simultaneously (Kaneko et al., 2004; Kovalchuk and Garaschuk, 2012; Gensch et al., 2015). Chloride indicator dyes, such as (6-Methoxyquinolinio) acetic acid ethyl ester bromide (MQAE) and its analogs, have been used to measure chloride levels *in vitro* (Verkman et al., 1989; Koncz and Daugirdas, 1994; Kovalchuk and Garaschuk, 2012; Gensch et al., 2015).

In this study, we utilized two-photon fluorescence lifetime imaging microscopy (2P-FLIM) to investigate activity dependent alterations in [Cl⁻]_in_ in dentate granule cells within organotypic hippocampal slice cultures. Specifically, we examined whether the application of the convulsant, pilocarpine, a non-specific muscarinic acetylcholine receptor agonist, could elevate neuronal activity and induce an increase in [Cl⁻]_in_ in these cells. It has been shown that excessive neuronal activity promotes KCC2 protein dephosphorylation, resulting in decreased KCC2 functionality (Lee et al., 2010). Therefore, investigating whether prolonged neuronal activity elevates intracellular chloride levels is crucial. By employing multicellular FLIM imaging and a Gaussian mixture statistical model (Takano et al., 2012), we analyzed the population distribution of intracellular chloride. Our approach revealed the emergence of subpopulations of dentate granule cells with elevated [Cl⁻]_in_ following prolonged pilocarpine exposure, originating from a largely homogeneous population. We will discuss the potential implications of these emerging subpopulations in chronic epilepsy.

## 1 MATERIALS and METHODS

### 1.1 Organotypic hippocampal slice cultures

All procedures were performed following the protocol approved by the Institutional Animal Care and Use Committee (IACUC) at the Children’s Hospital of Philadelphia. C57BL/6J mice (Jackson Laboratory) postnatal age P3-P6 were anesthetized with isoflurane and rapidly decapitated. The brains were removed and placed in an ice-cold dissection medium of 25 mM HEPES buffer solution with 6% glucose. The hippocampi were dissected and sliced transversely into 400 um sections using a McIlwain tissue chopper. Two to three slices with well-defined hippocampal neuronal layers were placed onto each semi-porous membrane insert (0.4 um, Millipore) inside a 35 mm culture dish. The brain slices were cultured with growth media consisting of 45% neurobasal medium, 25% heat-inactivated horse serum, 25% HBSS, 10 mM HEPES, 2 mM glutamine, and 6.5 mg/ml glucose in a humidified incubator with 5% CO_2_ at 37 °C. The cultures were maintained for 2-3 weeks with medium changes every 2–3 days. For a subset of experiments, acute slices were prepared from adult mice (C57BL/6J background, 2-3 months) as described in our earlier work (Dengler et al., 2017).

### 1.2 Dye loading, GCaMP6 transfection, and pilocarpine application

Organotypic slices were cultured for three weeks before being used for imaging experiments. To measure intracellular chloride levels, we used the membrane permeant chloride sensing dye, N-(Ethoxycarbonylmethyl)-6-methoxyquinolinium bromide (MQAE). The relative advantages of using MQAE are discussed in the Discussion section. The slice cultures were incubated with 10 mM MQAE in HEPES buffer solution for 20 minutes, washed three times, and imaged in a recording solution containing 155 mM NaCl, 3 mM KCl, 1 mM MgCl2, 3 mM CaCl2, 25 mM glucose, and 10 mM HEPES at a pH of 7.35. For prolonged pilocarpine application experiments, slices were treated with culture media containing pilocarpine (5 mM) for 24 hours, washed with standard culture media, and further incubated for 1-2 days before imaging. To perform calcium imaging, AAV-GCaMP6 viral particles were added to the slices after two weeks of culture and incubated for one week before imaging. In the third week, we imaged either control slices (no treatment) or slices treated with 5 mM pilocarpine.

### 1.3 Two-photon excitation fluorescence lifetime imaging (2P-FLIM)

A Becker-Hickle time-correlated photon counting system, a GaAsP PMT (Hamamatsu Photonics, H7422P40), and a power supply (Hamamatsu) were integrated onto a Prairie Technologies Ultima multiphoton microscopy system equipped with an Olympus BX61 microscope and a Ti: Sapphire femtosecond laser (Spectra Physics MaiTai HP DeepSee) for the imaging experiments. The MaiTai laser, with an 80 MHz pulse repetition rate, was used to trigger time-correlated photon counting. MQAE fluorescence was excited at 780 nm, and emission was collected with an emission filter of 470/30 nm. Photons were collected at less than half a million photon counts per second, and the photon counts were integrated for 120 seconds to capture a single data set. The resulting data was a three-dimensional dataset with 256 x 256 pixels and 256 time points. The Becker & Hickle SPI image software was used to determine fluorescence lifetime using a curve-fitting method with incomplete decay. The total number of photons detected per pixel provided an “intensity” value. The lifetime and intensity maps were exported as text images from the SPI software. For live/dead cell assays, a Thorlabs Bergamo Scope with a Ti:Sapphire femtosecond laser (Spectra Physics MaiTai HP DeepSee) was utilized. In this study, we will utilize fluorescence lifetime in nanoseconds as a relative measure of Cl-levels. The use of time units to represent chloride levels is a common practice in MQAE FLIM studies (Gagnon et al., 2013; Untiet et al., 2017).

### 1.4 Image and statistical analysis

Using ImageJ, circular or oval regions of interest (ROIs) were manually drawn over the cell body area in the image generated by total photon counts. The ROIs were then transferred to the corresponding lifetime image, and each cell’s average MQAE fluorescence lifetime within the ROI was calculated. A model-based cluster analysis, i.e., the Gaussian mixture model, was used to categorize cells based on the lifetime distribution. This is a parametric statistical method for clustering data into several unobserved classes (Takano et al., 2012). The modeling process involves selecting the number of classes based on the minimum quantity that maximizes the Bayesian Information Criteria (BIC) metric. The model then estimates the mean and standard deviation for each class and assigns a posterior probability of membership to each cell. Cells are assigned to a class based on their maximum posterior probability. The analysis was conducted using the mclust library (version 6.1.1,) in R4.4.1 (R Foundation for Statistical Computing).

## 2 RESULTS

### 2.1 MQAE 2P-FLIM of Dentate Granule Cells in hippocampal organotypic slices

First, we verified that MQAE 2P-FLIM can effectively report intracellular chloride levels in a quantitative manner by using calibration solutions with known chloride concentrations. Since the fluorescence lifetime of MQAE can be affected by various ionic species, such as proteins and peptides (Kaneko et al., 2004), it is important to calibrate the MQAE lifetime in the same cell types under investigation. Figure 1 shows an MQAE fluorescence intensity map and color-coded fluorescence lifetime in dentate granule cells (DGCs) in hippocampal organotypic slice cultures incubated in known extracellular chloride concentrations, along with the chloride-ionophore tributyltin-chloride, and the proton/potassium exchanger nigericin (Gensch et al., 2015). The intensity maps in the top row illustrate that higher chloride concentrations result in dimmer fluorescence, consistent with the notion that chloride ions quench MQAE fluorescence. Although fluorescence intensity has been used to measure relative changes in chloride levels (Koncz and Daugirdas, 1994), an intensity-based method can be affected by variations in dye loading. As fluorescence lifetime is less affected by the amount of dyes loaded in the cell, the FLIM approach offers advantages (Gensch et al., 2015; Untiet et al., 2017). Fig.1C shows the inverse of mean lifetime values plotted as a function of chloride levels. The linear relationship is consistent with MQAE fluorescence being quenched by chloride ions through a collisional quenching process. The Stern-Volmer constant (K_sv_) was estimated to be 45.8 M^−1^, by normalizing the data with an estimated y-intercept value, τ_0_ of 9.49 ns. In this study, we utilize fluorescence lifetime (τ) in nanoseconds or reciprocal of lifetime (1/τ) as a relative measure of Cl^−^ levels. The use of time (or inverse of time) to represent relative chloride levels or change in chloride levels is common practice in MQAE FLIM studies (Gagnon et al., 2013; Untiet et al., 2017).

**Figure 1.**
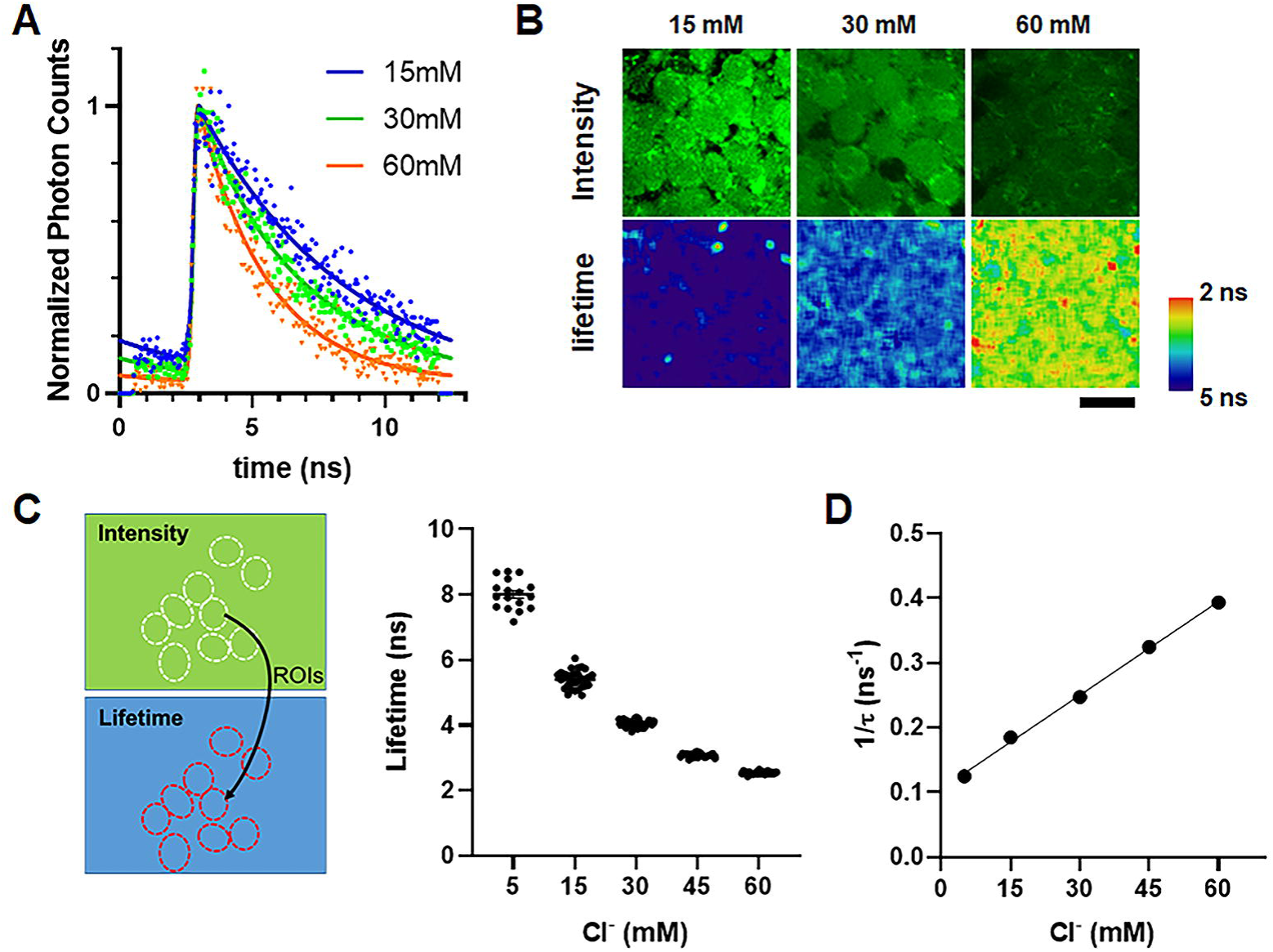
MQAE calibration in dentate granule cells in hippocampal organotypic slice culture. We imaged dentate granule cells (DGCs) in calibration solutions containing various chloride concentrations: potassium chloride (KCl) (5, 15, 30, 45, and 60 mM), potassium nitrate (KNO3)(145, 135, 120, 105, and 90 mM, respectively), HEPES (10 mM), 10 mM glucose (10 mM), tributyltin chloride (an ionophore for chloride ions, 10 μM), and nigericin (a proton/potassium exchanger, 10 μM), maintaining a pH of 7.35 adjusted with potassium hydroxide. **(A)** Each image pixel consists of time-correlated photon counts. The plot shows normalized photon counts obtained from three different slices/conditions. Higher chloride levels in the calibration solution correspond to faster fluorescence decay. Fluorescence lifetime was determined from the decay curves for each pixel in the 256×256 imaging frame. **(B)** (Top) Fluorescence intensity maps indicated by the total photon counts per pixel. Individual cell bodies are clearly identifiable. (Bottom) Fluorescence lifetime maps are color-coded, with warmer colors indicating higher chloride levels. Scale bar = 20 μm. **(C)** (Left) Regions of interest (ROIs) were manually drawn on the neuronal cell bodies in the intensity maps and then transferred to the corresponding lifetime maps. (Right) The average fluorescence lifetime was plotted as a function of chloride levels in the calibration solutions (black circles). **(D)** Inverse of lifetime values were plotted as a function of chloride concentration of the calibration solution. The linear relationship is consistent with MQAE fluorescence being quenched by chloride ions through collisional quenching process.

### 2.2 Application of furosemide, a KCC2 blocker, induced an increase in chloride levels in DG cells

First, we assessed baseline intracellular chloride levels in dentate granule cells in organotypic slice cultures. Figure 2A (top, left) displays the fluorescence intensity map, or total photon counts of MQAE-stained DG cells, imaged approximately 20 μm below the culture surface. Individual cells in the granule cell layer (GCL) are discernible. Approximately 70 cell bodies were identified in the GCL within this field of view. Figure 2A (bottom, left) shows the corresponding reciprocal of fluorescence lifetime (1/τ), where warmer colors indicate higher chloride levels. Using a circular or oval region of interest (ROI), we measured the average 1/τ over the cell body. The histogram in Figure 2B displays the distribution of the 1/τ, centered around 0.22 ns^−1^.

**Figure 2.**
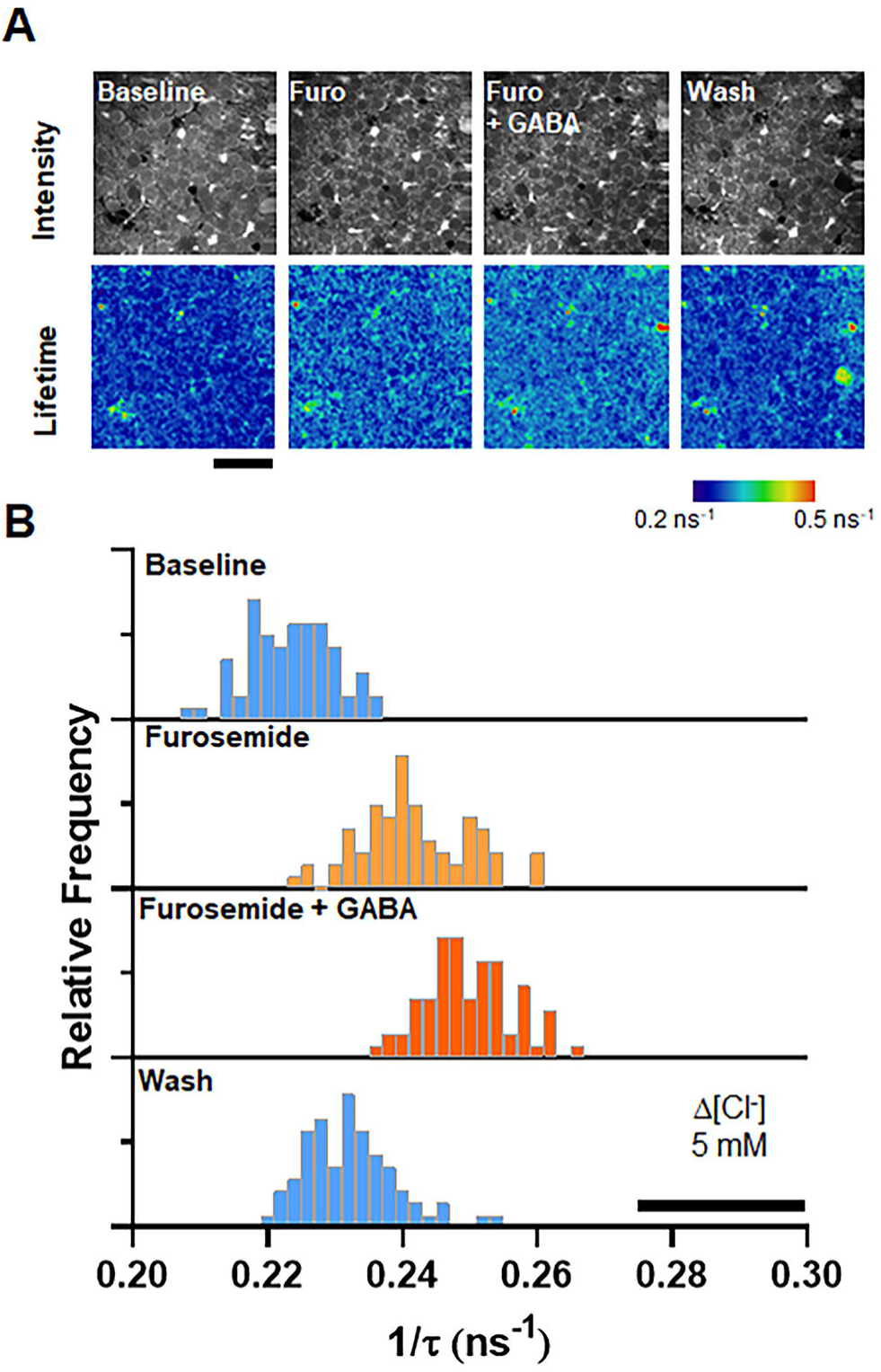
Application of furosemide, a KCC2 blocker, induced an increase in chloride levels in DG cells. **(A)** We assessed baseline intracellular chloride levels in dentate granule cells in organotypic slice cultures. The image on the top/left displays the fluorescence intensity map of MQAE-stained DG cells. Approximately 70 cell bodies were identified in the GCL within this field of view. The image on the bottom / left shows the corresponding reciprocal of fluorescence lifetime (1/τ), where warmer colors indicate higher chloride levels. Scale bar = 50 μm. Using a circular or oval region of interest (ROI), we measured the average 1/τ over the cell body. **(B)** The histogram in B displays the distribution of the reciprocal of fluorescence lifetime, centered around 0.22 ns^−1^. Application of an imaging solution containing 1mM furosemide, a KCC2 blocker, elevated chloride levels by ∼ 5 mM. The addition of 100μM GABA further increased the chloride levels, implying the activation of GABA receptors promoted chloride ion influx and accumulation. The elevated chloride level was reversed to baseline level after washing with a normal imaging solution.

To evaluate the effectiveness of the chloride extruder KCC2 function, we applied an imaging solution containing 1 mM furosemide, an antagonist of both KCC2 and NKCC1. We predicted that inhibiting KCC2 would lead to chloride ion accumulation, resulting in shortened MQAE lifetime, or increase in 1/τ values. Figure 2A (second from left) displays color-coded 1/τ map during furosemide application. The histogram in Fig.2B shows 1/τ increased to ∼0.24 ns^−1^, indicating that blocking KCC2 activity caused a few mM increase in [Cl^−^]_in_. The addition of 10 μM GABA to activate GABA_A_ receptors, promoting chloride ion entry, further increased 1/τ to ∼0.25 ns^−1^, corresponding to an overall [Cl^−^]_in_ increase of ∼5 mM. Washing the slice with regular imaging solution reduced 1/τ back to nearly the baseline of 0.23 ns^−1^. Overall, this demonstrates that we can induce changes in chloride levels and effectively detect the relative shift of [Cl^−^]_in_ using MQAE FLIM approach.

### 2.3 Acute Pilocarpine Application Induces Robust Synchronous Activity and Alters Intracellular Chloride Levels in Subpopulation of Neurons

Pilocarpine is a non-specific agonist of muscarinic acetylcholine receptors (mAChRs), which activates neurons via G-protein-coupled downstream pathways, primarily decreasing activity of GABAergic interneurons. Since mAChRs are abundant in the dentate gyrus, CA3, and CA1 regions, pilocarpine enhances neuronal activity throughout the hippocampus. To assess the effects of pilocarpine application on neuronal activity, we conducted calcium imaging using slice cultures expressing GCaMP6. Under normal imaging conditions (HEPES-buffered aCSF) without pilocarpine, dentate granule neurons exhibited minimal spontaneous firing activity. In Figure 3A, the top displays the baseline fluorescence, and the bottom shows a snapshot of the normalized fluorescence change (ΔFmax/F_0_) at the peak timing indicated in Figure 3B. Figure 3B displays the time course of the normalized fluorescence change (ΔFmax/F_0_) averaged across the entire field of view. The average peak height for the spontaneous activity was approximately 5% (ΔFmax/F0 = 0.05). This activity was mainly localized in neuronal processes surrounding the cell bodies, with only a few cells responding in the cell body region (Figure 2A, bottom). In the imaging solution containing 5 mM pilocarpine (Poulsen et al., 2002), neurons exhibited highly intense, synchronous spontaneous firing activity. Figures 3C and 3D show responses from numerous cell bodies visible as donut-like shapes (Figure 3C, bottom), and the maximum response reached a 120% increase (ΔFmax/F0 = 1.2) (Figure 3D). These activities are consistent with the firing of multiple action potentials, as indicated by the calcium transient amplitude (Takano et al., 2012; Yu et al., 2013; Kuzum et al., 2014). The frequency of large synchronous network activity was approximately 0.03 Hz, with an average inter-event interval of 30 seconds.

**Figure 3.**
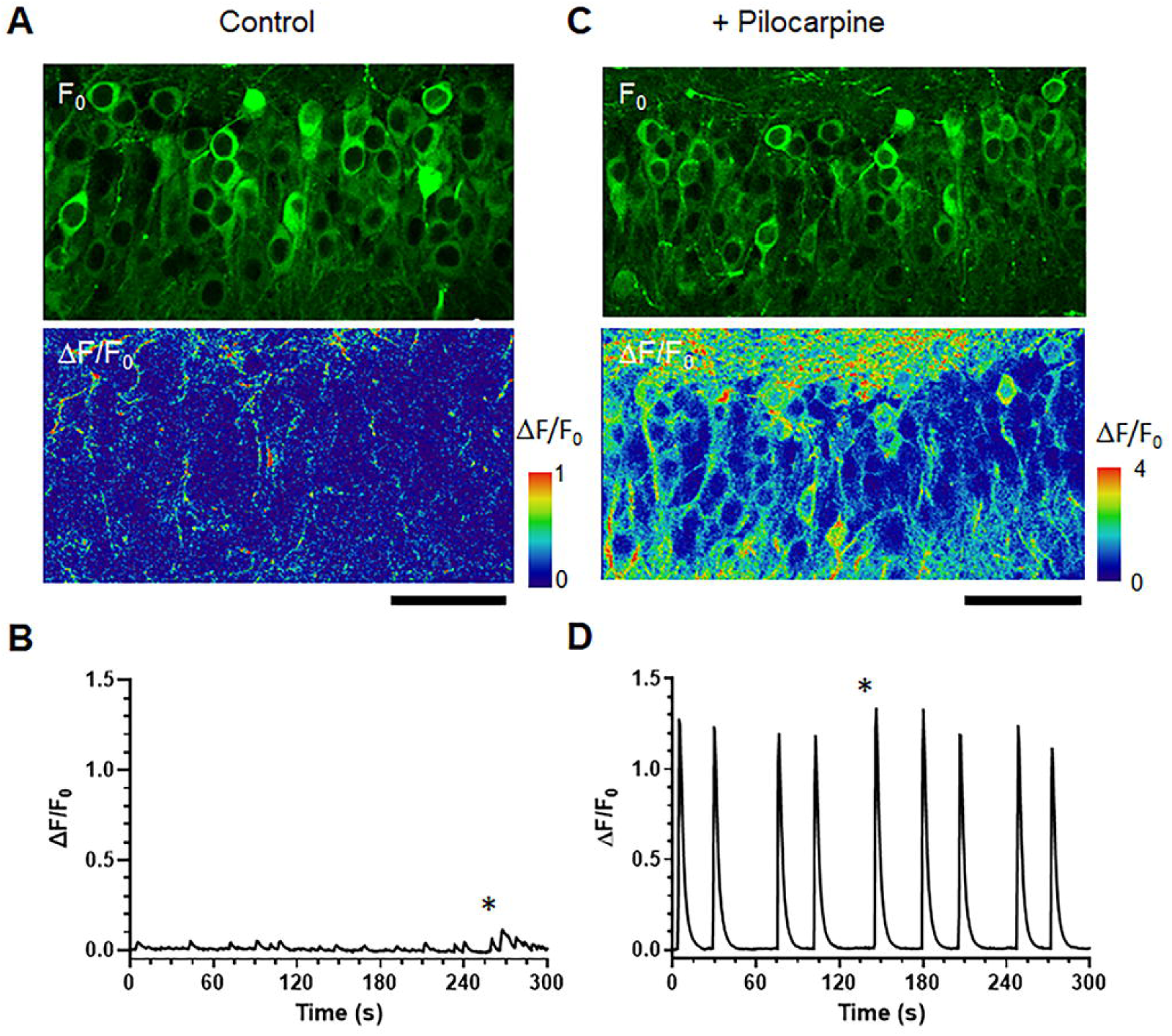
Pilocarpine induces robust spontaneous synchronous activity in the slice culture. **(A)** Time-series fluorescence imaging of dentate granule cells expressing GCaMP6 was captured under control conditions. (Top) Baseline fluorescence (F_0_). (Bottom) Normalized fluorescence change (ΔF/F_0_) at the peak timing indicated by the star in B. **(B)** Average ΔF/F_0_ over time over across the entire field shown in A. The dentate granule cells exhibited slight but visible spontaneous activity in the control condition. Responses in neuronal processes are noticeable, but less obvious in cell bodies. Scale bar = 50 μm. **(C)** Time-series fluorescence imaging of dentate granule cells in an imaging solution containing 5mM pilocarpine. (Top) Baseline fluorescence. (Bottom) ΔF/F_0_ at the activity peak indicated by the star in D. **(D)** Average ΔF/F_0_ trace over the entire field shown in C. Dentate granule cells exhibited robust spontaneous activities in both neuronal processes and in the cell bodies. Judging from the amplitude at the peak (ΔF/F_0_ > 1) and duration of the calcium transients (∼10 seconds), as well as the fact that all cells within the field of view fired simultaneously, the calcium transients consist of network bursting events. The continuous network bursting mimics the status epilepticus in animal model of epilepsy, induced by systemic injection of pilocarpine.

To assess the acute effects of pilocarpine-induced intense synchronous activity on intracellular Cl^−^ levels, we performed a time-series MQAE FLIM analysis. Figure 4A shows intensity and fluorescence lifetime images (1/τ) captured at 0 to 20 minutes during exposure to 5 mM pilocarpine. The color coded 1/τ images display warmer colors after 20 minutes. Figure 4B presents the mean 1/τ values measured over six randomly selected cell body regions (ROIs). Some cells exhibited a gradual increase in 1/τ values, while others maintained unchanged 1/τ values. The histogram of 1/τ in Figure 4C shows a small shift in the mean value from 0.24 ns^−1^ to 0.25 ns^−1^, with several cells shifting to larger values up to ∼0.32 ns^−1^. The overall shift in chloride levels may be reaching ∼5 mM in some cells.

**Figure 4.**
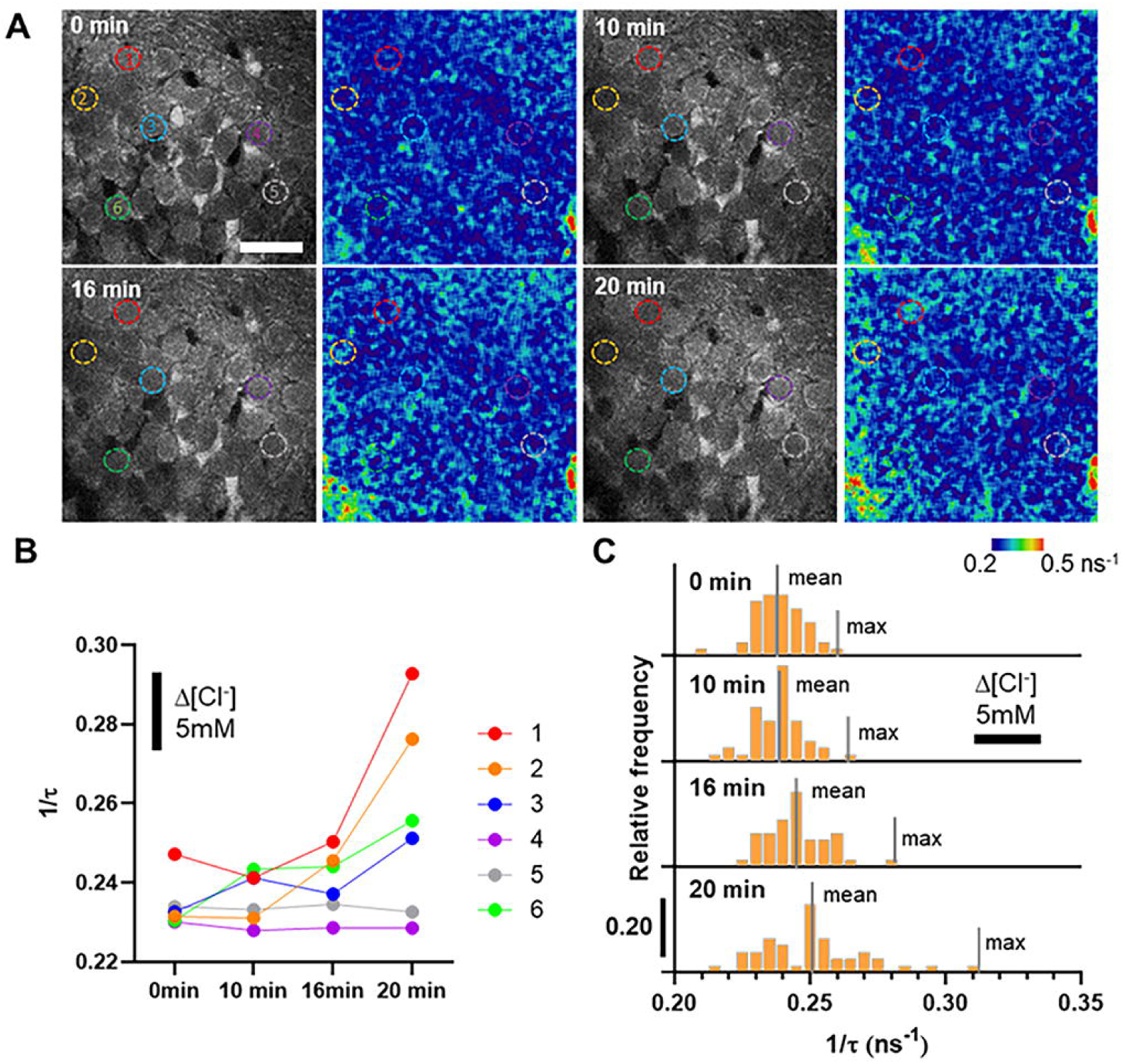
Acute Pilocarpine Application Moderately Increases Intracellular Chloride Levels. **(A)** Time-series FLIM of dentate granule cells in an imaging solution containing 5 mM pilocarpine. The intensity map (left) and the color-coded reciprocal of fluorescence lifetimes (1/τ) (right) at 0 min, 10min, 16min, 20 min after pilocarpine application are shown. An increased presence of warmer colors in the 1/τ images at 16 and 20 minutes suggests a slight increase in chloride levels. Scale bar = 30 μm. **(B)** 1/τ values of six randomly selected cells (ROIs) in (A), showing a gradual increase in 1/τ values for some cells (cells 1, 2, 3, and 6), while 1/τ values remained constant for others (cells 4 and 5). **(C)** Histogram of 1/τ for all cells in the images in (A), with the population mean and maximum values indicated by lines. The 1/τ values gradually shift to higher values. Based on the scale bar corresponding to the relative change in chloride levels, the chloride level may have increased by approximately 5 mM for some cells.

### 2.4 Prolonged Pilocarpine Exposure Induces Massive Increases in Intracellular Chloride Levels

In a previous study of pilocarpine-induced seizure-like activity in hippocampal slice cultures (Poulsen et al., 2002), exposure to 5 mM pilocarpine for 24 hours significantly increased brain-derived neurotrophic factor (BDNF) expression. Elevated BDNF levels have been observed in both epilepsy patients and *in vivo* models of epilepsy. Therefore, the study authors suggested that prolonged pilocarpine exposure in hippocampal slice cultures could serve as an *in vitro* model for epilepsy. To replicate this experimental protocol, we added 5 mM pilocarpine to the culture media and incubated for 24 hours. Subsequently, we washed the slices and cultured them in standard culture media for 1-2 days.

Figure 5 compares MQAE FLIM images under control conditions and after 24 hours of pilocarpine exposures. Fig.5A (left) shows a representative fluorescence intensity map of DG cells, imaged approximately 20 μm below the culture surface. Individual cells in the granule cell layer (GCL) are evident, and the hilus can be distinguished from the GCL. Approximately 40 cell bodies were identified in the GCL within this field of view. Variability in fluorescence intensity was observed in the soma, as seen in the darker cell marked by the blue arrow and the darkest cell marked by the red arrow. Figure 5A (right) shows the corresponding reciprocal of fluorescence lifetime (1/τ), where warmer colors indicate higher chloride levels. Using a circular/oval region of interest (ROI), we measured average 1/τ over the cell body region. Under control conditions, we evaluated 11 slices prepared from 8 animals. In Fig.5C (top), the histogram of the inverse of lifetime for all cells (n = 571 cells (ROIs) is plotted as relative frequency with a peak at ∼0.237 ns^−1^ and a scale bar indicating the relative difference in chloride levels.

**Figure 5.**
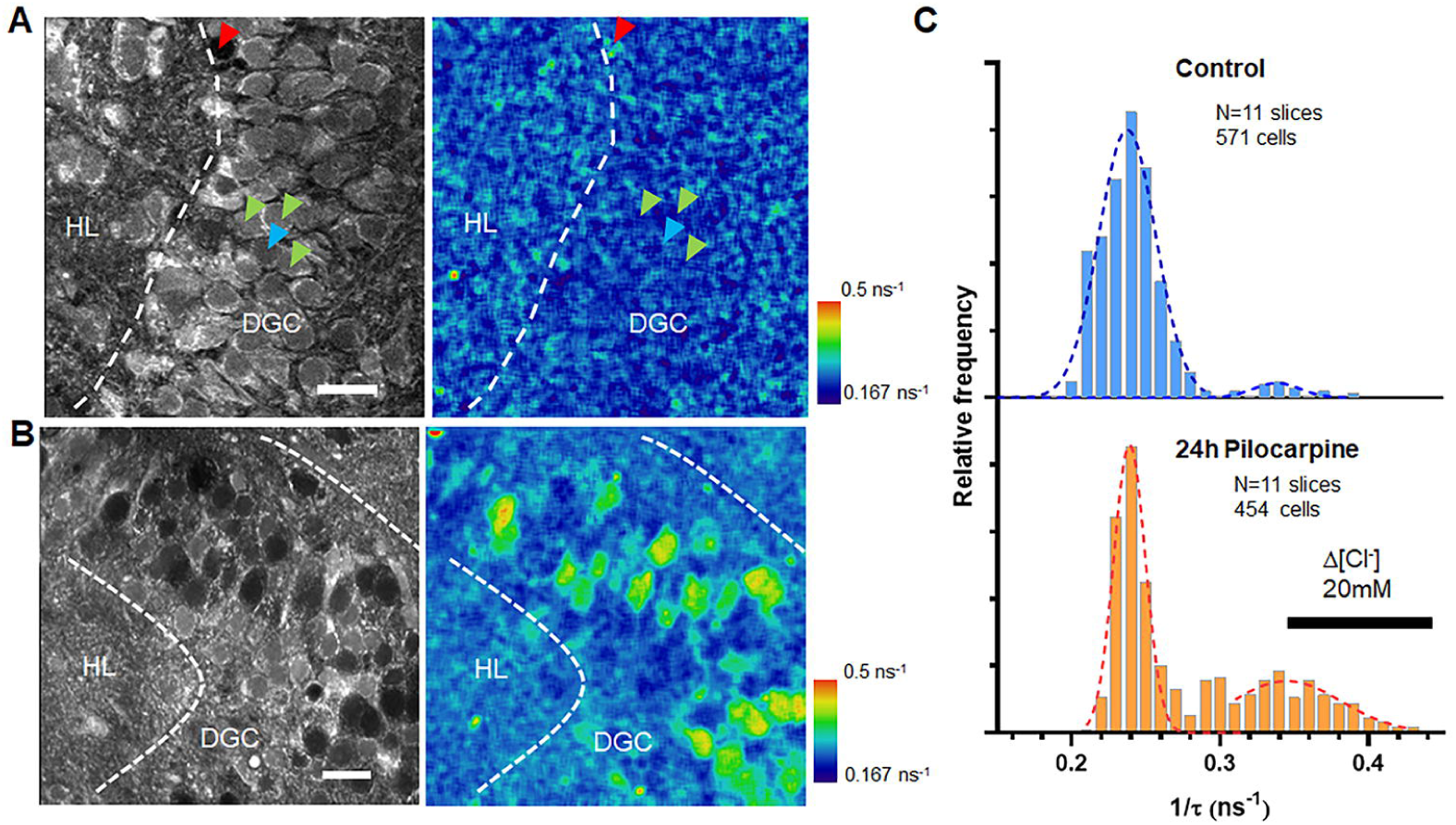
Prolonged Pilocarpine Exposure Induces Massive Increases in Intracellular Chloride Levels in a Subpopulation of Neurons.. **(A)** A fluorescent intensity map and corresponding reciprocal of fluorescence lifetime (1/τ) of dentate granule cells (DGC) in a hippocampal slice culture imaged 20 microns below the surface. **(B)** A fluorescent intensity map and corresponding reciprocal of 1/τ of DGC in a hippocampal slice culture incubated with pilocarpine for 24 hours prior to imaging in a normal imaging solution (details provided in the method section). In the granule cell layer, the dark circular regions coexist with the lighter-colored circular shapes, similar to those observed in the control condition. The dark regions seen in the intensity map exhibit much larger 1/τ values. **(C)** A histogram of 1/τ values for 571 cells (ROIs) from 11 slices prepared under the control shows a peak around 0.23 ns^−1^. Another histogram of 1/τ obtained from 454 cells (ROIs) from 11 slices incubated with 5mM pilocarpine for 24 hours shows a bimodal distribution, with one peak around 0.23 ns and another peak around 0.34 ns^−1^. This indicates that the pilocarpine treatment induced the emergence of a subpopulation with higher chloride levels. Scale bars = 20 μm.

Fig.5B shows a representative image from the slices treated with pilocarpine for 24 hours. The main difference from the control slice is the presence of darker regions in the cell body layer. The corresponding 1/τ image reveals areas with shorter lifetimes (indicated by warmer colors). We evaluated 11 slices from 7 animals 1-2 days following pilocarpine treatment. The histogram of the reciprocal of lifetimes for all cells (n = 454 cells (ROIs)) shows a bimodal distribution, with one peak at around 0.239 ns^−1^and the other around 0.344 ns^−1^. The corresponding difference in chloride levels is estimated to be ∼20 mM. The peak of the lower chloride population closely matched that of the control condition in Fig. 5C (top). Roughly half of the neurons maintained a similar chloride level to that of the control slices. A nonparametric Mann-Whitney test between the control and the pilocarpine-treated slices showed a significant difference (p<0.001).

### 2.5 Analysis of Multicellular Imaging Data using Gaussian Mixture Model

To quantify the multiple populations of varying [Cl^−^]_in_ in DGCs in a more rigorous manner, we utilized Gaussian mixture model-based cluster analysis (Takano et al., 2012) to analyze the dataset presented in Figure 5. For each slice preparation, with the number of cells ranging from 33 to 60, we conducted model-based clustering analysis using the statistical software R and its *mclust* library to fit the lifetime data to multiple Gaussian scenarios, ranging from one to five Gaussian distributions, either with equal variance or unequal variance. We calculated the Bayesian Information Criteria (BIC) for each scenario to determine the best fit for the slice preparation. As an example, for a pilocarpine exposed slice, the dataset was best fit by a model with two Gaussians of unequal variance, as indicated by the maximum BIC for this scenario (Fig6A). The model-based clustering method assigned a posterior probability of membership in each group (Gaussian) to each cell, with the highest posterior probability determining the group membership for each cell. Figure 6B depicts a histogram of the data, color-coded based on the assigned groups. For this slice, the mean values of the two Gaussians were 3.0 ns and 3.8 ns, respectively. Note that we used lifetime values rather than the reciprocal of lifetime for this analysis, as the raw lifetime values fit better with the Gaussian distribution.

**Figure 6.**
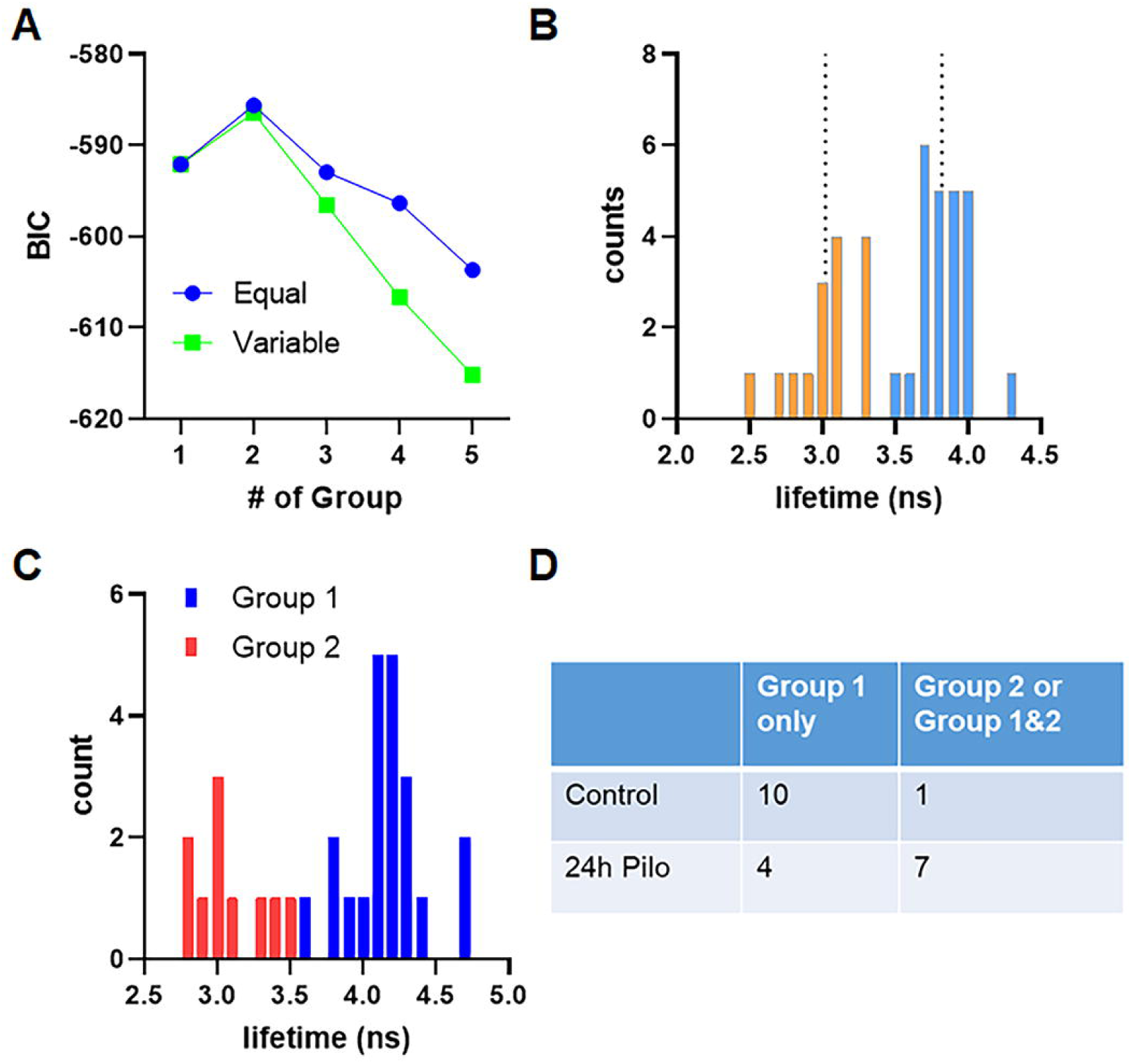
Gaussian mixture model-based cluster analysis for multicellular chloride imaging data. To objectively categorize cells into groups based on chloride levels, the fluorescence lifetime data were analyzed by model-based clustering with the *mclust* package in the R statistical analysis software. The analysis examined up to five Gaussian distributions with either equal variance or unequal variance. For each scenarios, Bayesian Information Criteria (BIC) were calculated. **(A)** The maximum BIC defines the best-fit model among the tested scenarios. In this data set (a pilocarpine-treated slice), the best-fit model was two Gaussians with equal variance. **(B)** Based on the best-fit scenario in A, probabilities of membership in each group were calculated for each cell. The highest probability determined the membership assignment for each cell. The plot shows a color-coded histogram reflecting the membership assignment, with mean values for each group showing in lines. **(C)** For both control and pilocarpine-treated slices, we identified a total of 33 sub-groups for 22 slices. By conducting the *mclust* analysis on the mean values of the sub-groups (33 values), we identified two major cgroups that has longer lifetime (Group1) and shorter lifetime (Group2). **(D)** We made a contingency table base on the grouping types (Group1 only vs Group2 or Group 1&2) the each slice belongs to. <u>Fisher’s exact test</u> shows significant difference between control condition and the pilocarpine-treated samples (p=0.0237).

We conducted the model-based clustering analysis for each slice under both control and pilocarpine exposed conditions (11 slices for each condition, total of 22 slices). Overall, we identified one to three Gaussian distributions for each slice, and a total of 33 Gaussians with mean values ranging from 2.78 ns to 4.67 ns. To ensue unbiased judgement, we further conducted model based cluster analysis on these mean values (n=33) and identified two groups as the best fit: Group 1 with a mean value 4.16 ns and Group 2 with a mean value of 3.08 ns. Under control conditions, most of the slices (10 out of 11) had a distribution belong only to Group 1, with one slice exhibiting two Gaussians from Group 1 and Group 2. In contract, for the pilocarpine treated slices, the majority (7 out of 11) had a Gaussian from Group 2, with five slices exhibiting multiple Gaussians from both Group 1 and Group 2. Fig.6D shows a contingency table describing these observations. A chi-square test revealed that the pilocarpine-exposed condition significantly differed from the control condition in the involvement of Group 2 (p=0.027). In other words, this supports the idea of the emergence of Group 2 population when the slice is exposed to pilocarpine for an extended period of time (24 hours).

### 2.6 Prolonged Pilocarpine Exposure did not cause significant Cell Death in DGCs

We then conducted a cell viability assay using a fluorescence probe that explicitly stains the nuclei of dead cells to investigate whether the increased chloride levels in pilocarpine-treated tissues are associated with cell death. Representative images of control and pilocarpine-treated slice cultures are presented in Figure 7. We found no significant difference in the number of dead cells in the DGC layers (indicated by red fluorescence) between the two conditions, despite a presumably high intracellular chloride population in about half of DGCs (see Figs. 5-6). In the hilus region, however, we observed more dead cells in the pilocarpine-treated slices than in the control condition. This observation is consistent with the in vivo pilocarpine epilepsy model, where significant interneuron loss in the hilus region has been reported (Buckmaster et al., 2017).

**Figure 7.**
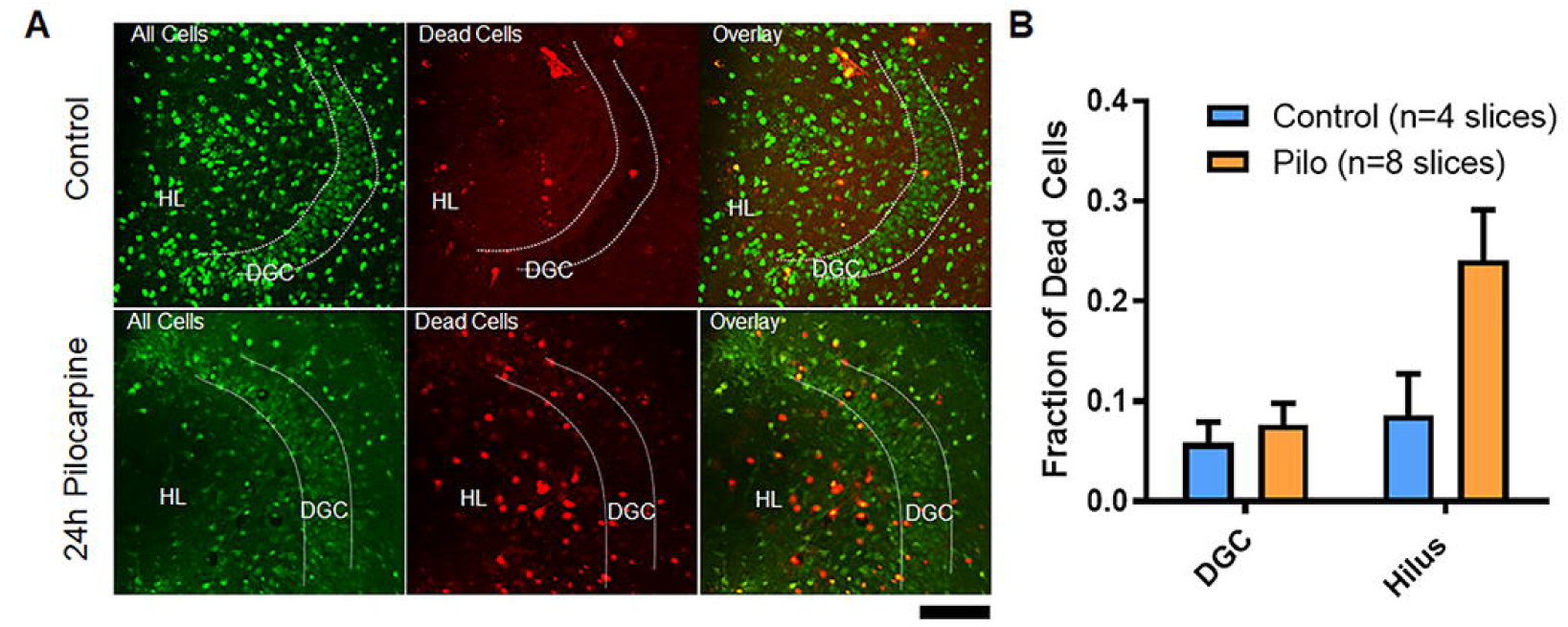
Live/dead cell assay. To determine whether the elevated chloride levels resulting from prolonged pilocarpine exposure led to cell death, we conducted a fluorescence-based live/dead cell assay. A green fluorescence dye stains all cells, regardless of viability, while a red fluorescence probe stains the nuclei of dead cells. **(A)** Using conventional two-photon microscopy, we captured green/red fluorescence micrographs. The top row shows example images of dentate granule cells from a slice culture prepared under the control condition, while the bottom row presents example images of a slice culture incubated with pilocarpine for 24 hours. In both cases, green fluorescence cells are clearly visible in dentate granule cell layer, and the number of red fluorescence cells is comparable. However, in pilocarpine treated slice, more red fluorescence cells exists in hilus region. Scale bar = 100 μm. **(B)** The number of cells (green) and dead cells (red) in each regions (granule cell layer vs. hilus) were manually counted using imageJ software and quantified across multiple slice culture preparations. The faction of dead cells did not exhibit significant differences between the two conditions, but the fraction of dead cells in the hilus was significantly higher in pilocarpine treated slices (Two-way ANOVA p=0.03; Bonferroni’s multiple comparisons: Control vs Pilo - DGC not significant, Hilus significant).

## 3 DISCUSSION

### 3.1 Emergence of Bimodal Population During Status Epilepticus

In a previous electrophysiological study using rats, gramicidin perforated patch recordings of DGC neurons revealed a subpopulation of neurons with depolarized GABA reversal potential in the slices prepared from rats 7 days after pilocarpine-induced status epileptics (Pathak et al., 2007). The data suggest a bimodal distribution of DGC neurons with one population exhibiting a normal GABA reversal potential with approximately at −70 mV and and a second population exhibiting a depolarized GABA reversal potential with a peak around −50 mV. Intracellular chloride levels can be estimated by assuming a parameter like permeability ratio (α) of carbonate ions and chloride ions (Berglund et al., 2006). With α of 0.3, the shift in GABA reversal potential from −70 mV to −50 mV corresponds to chloride level increase of approximately 18 mM. In this study, we observed that acute pilocarpine application caused some cells to exhibit slightly elevated chloride levels (approximately 5 mM increase) within 20 minutes exposure (Fig.4). Prolonged exposure to pilocarpine in the culture media for 24 hours caused more dramatic changes; roughly half of cell had significantly elevated chloride levels (approximately a 20 mM increase), while the rest of cells maintained chloride levels similar to the control condition (Fig.5). This change was similar in magnitude to the E_GABA_ measurements in slices from the pilocarpine animal model discussed above.

Pilocarpine might have contributed to an activity-dependent reduction in KCC2 function through tyrosine phosphorylation, as demonstrated by Lee et al. (Lee et al., 2010). They showed that prolonged activation of mAChRs enhanced KCC2 tyrosine phosphorylation in cultured hippocampal neurons. This reduction in KCC2 function would have an immediate effect. Therefore, the experiment shown in Figure 4, where we observed a moderate increase in the chloride level in most cells within 20 minutes, could be linked to the KCC2 phosphorylation process within subpopulations of DGCs. However, the magnitude of changes observed through pilocarpine application, which was similar to the changes induced by blocking KCC2 with furosemide (Fig.2), was much smaller than the mean difference observed between the two Gaussians during the prolonged pilocarpine applications. These data suggest that the reduced chloride extrusion function may not be the sole cause of the increased chloride level.

The route of chloride ion influx remains unknown. It is less likely that the chloride accumulator Na-K-2Cl cotransporter isoform1 (NKCC1) is expressed to any significant extent in adult neurons. GABA_A_ receptor channels could be a source of chloride influx, as demonstrated in Fig.2, where the addition of GABA led to an elevation of chloride levels. GABA should promote chloride influx under normal conditions, and the repeated actvation of GABAergic interneurons could lead to chloride accumulation at a local site (Rahmati et al., 2021). Voltage-gated chloride channels or Cl^−^/HCO3^−^ exchangers could also contribute to chloride ion influx. A voltage-gated chloride channel, SLC26A11,was implicated in the cellular edema process (Rungta et al., 2015). Since the increase in chloride level after a 24-hour pilocarpine treatment was accompanied by an increase in cell size (data not shown), the prolonged pilocarpine exposure process might involve cellular edema. Cellular edema is a reversible process and cells are still alive, which is consistent with our live/dead cell assay (Fig.7) indicating that most cells are still viable in the DGC layer. Another critical function of KCC2, apart from being a chloride and potassium extruder, is acting as a water molecule extruder (Chamma et al., 2012). Thus, the reduced KCC2 function could cause water molecules to accumulate, resulting in increased cellular volume. This increase in cellular volume could help maintain chloride ion concentration at a normal level. This might be the case for some cells with large sizes but with a normal chloride level.

In the previous perforated patch study using epilepsy model rats (Pathak et al., 2007), the authors reported bimodal GABA reversal potentials one week after the status epileptics (SE), However, this phenomenon becomes normalized (single distribution) two weeks after SE and in chronic epileptic rats. This suggests that the elevated chloride levels are transient. Nonetheless, the authors noted that the normal chloride levels do not necessarily guarantee proper inhibitory function. In pathological conditions, a cell’s tolerance for chloride loading might change, leading to a rapid elevation of chloride levels with minimal GABA receptor activation. That could be the case for chronic epilepsy, where cells are easily excited (Dengler et al., 2017). These circumstances might result in a hyper-excitable dentate gyrus, which could lead to changes in behavioral outcomes (Kahn et al., 2019). If we can design a ‘chloride loading stress test’ using chloride imaging approaches, it could serve as a valuable assay tool for further delineating inhibitory functions.

### 3.2 Basal Chloride Levels in DGC Measured by MQAE FLIM Imaging

The chloride concentration reading from the FLIM experiment resulted in an apparent [Cl^−^]_in_ of approximately 26 mM in the baseline condition, which may be higher than the expected cytosolic chloride level in adult neurons. We expect the baseline chloride level to be as low as 10 mM to maintain proper inhibitory function of GABA_A_ receptors. Several scenarios may contribute to this observation. MQAE is membrane permeable and may penetrate cellular organelles such as liposomes and the endoplasmic reticulum (ER). It is known that organelle chloride levels can be as high as ∼100 mM (Hara-Chikuma et al., 2005; Chakraborty et al., 2017). In such cases, we may observe the overall average Cl^−^ levels, including those of the nucleus, organelles, and cytosolic components. Given that the cytosolic component is crucial for the proper functioning of GABA receptors, and that the cytosolic volume is relatively small compared to the nuclear volume in dentate granule cells, MQAE may not precisely reflect the chloride levels that directly determine GABA reversal potentials. Nevertheless, we are hopeful that the overall chloride concentration measured in the cell body (circular ROIs) positively correlates with the cytosolic levels that influence GABA receptor activity.

Note that slice cultures have been proposed as a model of trauma-induced epilepsy, where the initial slicing acts as a traumatic injury, and the subsequent slice culture exhibits spontaneous synchronous activity that mimics epileptic seizures (Lillis et al., 2015; Berdichevsky et al., 2016; Liu et al., 2017). Although the elevated chloride level (20-30 mM) in the control condition might be explained by this model, several observations suggest that the control condition still provides proper inhibitory function. These observations include: (1) the acute slices prepared from wildtype mice had similar MQAE lifetime (4.24 ± 0.26 ns, 0.237 ± 0.0154 ns^−1^, n = 424 cells (7 slices from 2 animals), data not shown); (2) the slice cultures in the control conditions did not have strong synchronous events in our hands (Fig.3); and (3) a clear difference in chloride levels was observed in pilocarpine-treated slices. Even if the chloride levels were indeed 20-30 mM and the action of GABA does not promote chloride ion influx, the increased conductivity resulting from GABA receptor channel opening (shunting effect) could still act to decrease excitability in control conditions. Due to this uncertainty, in this paper, we focused on the relative change in chloride levels between the control and pilocarpine-treated conditions by reporting MQAE lifetimes and reciprocal of the lifetime.

### 3.3 Multicellular Chloride Imaging

There are two major chloride imaging approaches available: the use of genetic probes such as Clomeleon (Berglund et al., 2006; Dzhala et al., 2010) and Superclomeleon (Rahmati et al., 2021), or the use of dyes such as MQAE (Gensch et al., 2015) and MQE (Zhang et al., 2006). Clomeleon and Superclomeleon are ratiometric probes based on chloride-sensitive YFP and chloride-insensitive CYP analogues and are extremely useful as measures of transmembrane chloride gradients. However, pH sensitivity is a disadvantage of these probes, which can be exacerbated by seizure-induced extracellular pH changes. The need for viral packaging and viral injection, and maintenance, as well as the availability of transgenic lines, are major disadvantages of genetic probes in tissue where viral transfection is not feasible. In contrast, despite the complexity in the interpretation of calibrated values discussed above, MQAE and its analogues have certain advantages. First, there is no need for genetic crossing or viral injection. They are not sensitive to pH, and their effective dissociation constant is in the physiological range. Although MQAE’s excitation wavelength is in the UV range with associated phototoxicity, employing a two-photon excitation approach eliminates this concern. Moreover, fluorescence lifetime imaging (FLIM) of MQAE is less affected by dye loading variation among tissues.

Chloride imaging has provided insights into the roles chloride ions play in various biological processes. For example, in an earlier study, MQE was used to measure chloride levels in retinal ganglion neurons in developing mice (Zhang et al., 2006), demonstrating the functional switch of GABA receptors from excitatory to inhibitory occurring around age P6. In the context of inhibitory function mediated by GABA receptors, chloride microdomains were studied using Super Cloemeleon as well as MQE (Rahmati et al., 2021). In a pioneering work on KCC2 function in pain circuitry (Gagnon et al., 2013), researchers used HEK cells expressing Clomeleon to screen for KCC2-enhancing small molecule drugs, and they found one (CLP257). They verified the chloride-extruding function using spinal slices and MQAE 2P-FLIM approach, showing that the application of CLP257 extended the MQAE lifetime, i.e., lowered the chloride levels. The application of chloride imaging goes beyond studying neuronal functions. Astrocytic chloride levels were studied, and increased chloride levels were observed when the KCC2 blocker, DIOA, was applied (Engels et al., 2021). In the most recent study, the role played by astrocytic chloride ions in synaptic transmission was studied (Untiet et al., 2024). They proposed that astrocytic chloride extrusion through GABA receptors on astrocytes provides modulatory effects on neuronal GABA synaptic transmission. They verified this by genetic probes expressed in astrocytes as well as by MQAE staining. Overall, classic dyes like MQAE can still provide useful information on cellular chloride regulation. They can be readily applicable to any tissues such as those from human patients or animals undergoing disease states. They can also be used in combination with genetic probes other than chloride sensors.

## 4. CONCLUSION

We investigated whether enhanced neuronal activity induced by pilocarpine application could lead to prolonged elevation in chloride ion concentrations. MQAE two-photon fluorescence lifetime imaging was performed on the control and pilocarpine-treated slices. Population analysis revealed that under control conditions, 95% of the cells belonged to a Gaussian distribution centered at 1/τ of 0.23 ns^−1^. The distribution shifted to a shorter lifetime in pilocarpine-treated slices, with nearly half of the population belonging to the distribution centered at 1/τ of 0.34 ns^−1^. In comparison, the remaining cells stayed in the group centered at 1/τ of 0.23 ns^−1^. These results suggest that the pilocarpine treatment increased [Cl^−^]_in_ in a subset of neurons. Our study demonstrated that the multicellular imaging approach combined with the Gaussian mixture model provided a powerful and rigorous method to categorize cells into groups in an unbiased and quantitative manner. We found that a subpopulation of neurons emerges from a previously homogeneous population upon application of pilocarpine. This phenomenon could be occurring following status epilepticus during the transition to an epileptic state.

## Acknowledgments

This work was supported by funding from the Children’s Hospital of Philadelphia Foerderer Grant (HT), the University of Pennsylvania Neuroscience Center grant (HT), and NIH NINDS R01NS038572 (DAC), R01NS082046 (DAC), the Intellectual and Developmental Disabilities Research Center (IDDRC) at CHOP/Penn NIH/NICHD P50HD105354.

## CRediT authorship contribution statement

**Hajime Takano:** Conceptualization, data curation, funding acquisition, data analysis, writing. **Fu-Chun Hsu:** Conceptualization, data curation, review & editing. **Srdjan Joksimovic**: data curation, review. **Doug A Coulter:** Conceptualization, funding acquisition, review & editing.

## Declaration of competing interest

The authors declare no competing financial interests.

## Notes

### Competing Interest Statement

The authors have declared no competing interest.

### Summary of Updates

One of the questions raised during review concerned why the control condition exhibited higher chloride levels than expected. Because all of the original experiments were conducted in slice cultures, this prompted the concern that results might differ in acute slices. To address this, we performed additional experiments in acute slices and measured chloride levels under control conditions. These results were consistent with those obtained in slice cultures, and the overall conclusions of the study remained unchanged. The manuscript was revised to incorporate these additional data, and we added as a co-author the colleague who performed the acute slice experiments.

## References

Allene C, Cattani A, Ackman JB, Bonifazi P, Aniksztejn L, Ben-Ari Y, Cossart R (2008) Sequential generation of two distinct synapse-driven network patterns in developing neocortex. J Neurosci 28:12851–12863.

Ben-Ari Y (2014) The GABA excitatory/inhibitory developmental sequence: a personal journey. Neuroscience 279:187–219.

Berdichevsky Y, Saponjian Y, Park KI, Roach B, Pouliot W, Lu K, Swiercz W, Dudek FE, Staley KJ (2016) Staged anticonvulsant screening for chronic epilepsy. Ann Clin Transl Neurol 3:908–923.

Berglund K, Schleich W, Krieger P, Loo LS, Wang D, Cant NB, Feng G, Augustine GJ, Kuner T (2006) Imaging synaptic inhibition in transgenic mice expressing the chloride indicator, Clomeleon. Brain Cell Biol 35:207–228.

Bonifazi P, Goldin M, Picardo MA, Jorquera I, Cattani A, Bianconi G, Represa A, Ben-Ari Y, Cossart R (2009) GABAergic hub neurons orchestrate synchrony in developing hippocampal networks. Science 326:1419–1424.

Buckmaster PS, Abrams E, Wen X (2017) Seizure frequency correlates with loss of dentate gyrus GABAergic neurons in a mouse model of temporal lobe epilepsy. J Comp Neurol 525:2592–2610.

Bui A, Kim HK, Maroso M, Soltesz I (2015) Microcircuits in Epilepsy: Heterogeneity and Hub Cells in Network Synchronization. Cold Spring Harb Perspect Med 5.

Cembrowski MS, N. (2019) Heterogeneity within classical cell types is the rule: lessons from hippocampal pyramidal neurons.. Nat Rev Neurosci 20:193–204.

Cess CG, Finley SD (2020) Data-driven analysis of a mechanistic model of CAR T cell signaling predicts effects of cell-to-cell heterogeneity. J Theor Biol 489:110125.

Chakraborty K, Leung K, Krishnan Y (2017) High lumenal chloride in the lysosome is critical for lysosome function. Elife 6.

Chamma I, Chevy Q, Poncer JC, Levi S (2012) Role of the neuronal K-Cl co-transporter KCC2 in inhibitory and excitatory neurotransmission. Front Cell Neurosci 6:5.

Cohen I, Navarro V, Clemenceau S, Baulac M, Miles R (2002) On the origin of interictal activity in human temporal lobe epilepsy in vitro. Science 298:1418–1421.

Dengler CG, Yue C, Takano H, Coulter DA (2017) Massively augmented hippocampal dentate granule cell activation accompanies epilepsy development. Sci Rep 7:42090.

Dzhala VI, Kuchibhotla KV, Glykys JC, Kahle KT, Swiercz WB, Feng G, Kuner T, Augustine GJ, Bacskai BJ, Staley KJ (2010) Progressive NKCC1-dependent neuronal chloride accumulation during neonatal seizures. J Neurosci 30:11745–11761.

Engels M, Kalia M, Rahmati S, Petersilie L, Kovermann P, van Putten M, Rose CR, Meijer HGE, Gensch T, Fahlke C (2021) Glial Chloride Homeostasis Under Transient Ischemic Stress. Front Cell Neurosci 15:735300.

Gagnon M, Bergeron MJ, Lavertu G, Castonguay A, Tripathy S, Bonin RP, Perez-Sanchez J, Boudreau D, Wang B, Dumas L, Valade I, Bachand K, Jacob-Wagner M, Tardif C, Kianicka I, Isenring P, Attardo G, Coull JA, De Koninck Y (2013) Chloride extrusion enhancers as novel therapeutics for neurological diseases. Nat Med 19:1524–1528.

Gensch T, Untiet V, Franzen A, Kovermann P, Fahlke C (2015) Determination of Intracellular Chloride Concentrations by Fluorescence Lifetime Imaging. In: Advanced Time-Correlated Single Photon Counting Applications (Becker W, ed), pp 189–211. Cham: Springer International Publishing.

Glykys J, Dzhala V, Egawa K, Balena T, Saponjian Y, Kuchibhotla KV, Bacskai BJ, Kahle KT, Zeuthen T, Staley KJ (2014) Local impermeant anions establish the neuronal chloride concentration. Science 343:670–675.

Hara-Chikuma M, Yang B, Sonawane ND, Sasaki S, Uchida S, Verkman AS (2005) ClC-3 chloride channels facilitate endosomal acidification and chloride accumulation. J Biol Chem 280:1241–1247.

Ikegaya Y, Aaron G, Cossart R, Aronov D, Lampl I, Ferster D, Yuste R (2004) Synfire chains and cortical songs: temporal modules of cortical activity. Science 304:559–564.

Kahle KT, Staley K (2008) Altered neuronal chloride homeostasis and excitatory GABAergic signaling in human temporal lobe epilepsy. Epilepsy Curr 8:51–53.

Kahn JB, Port RG, Yue C, Takano H, Coulter DA (2019) Circuit-based interventions in the dentate gyrus rescue epilepsy-associated cognitive dysfunction. Brain 142:2705–2721.

Kaneko H, Putzier I, Frings S, Kaupp UB, Gensch T (2004) Chloride accumulation in mammalian olfactory sensory neurons. J Neurosci 24:7931–7938.

Kelley MR, Cardarelli RA, Smalley JL, Ollerhead TA, Andrew PM, Brandon NJ, Deeb TZ, Moss SJ (2018) Locally Reducing KCC2 Activity in the Hippocampus is Sufficient to Induce Temporal Lobe Epilepsy. EBioMedicine 32:62–71.

Kinnunen PC, Luker KE, Luker GD, Linderman JJ (2021) Computational methods for characterizing and learning from heterogeneous cell signaling data. Curr Opin Syst Biol 26:98–108.

Koncz C, Daugirdas JT (1994) Use of MQAE for measurement of intracellular [Cl-] in cultured aortic smooth muscle cells. Am J Physiol 267:H2114–2123.

Kovalchuk Y, Garaschuk O (2012) Two-photon chloride imaging using MQAE in vitro and in vivo. Cold Spring Harb Protoc 2012:778–785.

Kuzum D, Takano H, Shim E, Reed JC, Juul H, Richardson AG, de Vries J, Bink H, Dichter MA, Lucas TH, Coulter DA, Cubukcu E, Litt B (2014) Transparent and flexible low noise graphene electrodes for simultaneous electrophysiology and neuroimaging. Nat Commun 5:5259.

Lee HH, Jurd R, Moss SJ (2010) Tyrosine phosphorylation regulates the membrane trafficking of the potassium chloride co-transporter KCC2. Mol Cell Neurosci 45:173–179.

Lillis KP, Wang Z, Mail M, Zhao GQ, Berdichevsky Y, Bacskai B, Staley KJ (2015) Evolution of Network Synchronization during Early Epileptogenesis Parallels Synaptic Circuit Alterations. J Neurosci 35:9920–9934.

Liu J, Saponjian Y, Mahoney MM, Staley KJ, Berdichevsky Y (2017) Epileptogenesis in organotypic hippocampal cultures has limited dependence on culture medium composition. PLoS One 12:e0172677.

Pathak HR, Weissinger F, Terunuma M, Carlson GC, Hsu FC, Moss SJ, Coulter DA (2007) Disrupted dentate granule cell chloride regulation enhances synaptic excitability during development of temporal lobe epilepsy. J Neurosci 27:14012–14022.

Poulsen FR, Jahnsen H, Blaabjerg M, Zimmer J (2002) Pilocarpine-induced seizure-like activity with increased BNDF and neuropeptide Y expression in organotypic hippocampal slice cultures. Brain Res 950:103–118.

Rahmati N, Normoyle KP, Glykys J, Dzhala VI, Lillis KP, Kahle KT, Raiyyani R, Jacob T, Staley KJ (2021) Unique Actions of GABA Arising from Cytoplasmic Chloride Microdomains. J Neurosci 41:4957–4975.

Rungta RL, Choi HB, Tyson JR, Malik A, Dissing-Olesen L, Lin PJC, Cain SM, Cullis PR, Snutch TP, MacVicar BA (2015) The cellular mechanisms of neuronal swelling underlying cytotoxic edema. Cell 161:610–621.

Takano H, McCartney M, Ortinski PI, Yue C, Putt ME, Coulter DA (2012) Deterministic and stochastic neuronal contributions to distinct synchronous CA3 network bursts. J Neurosci 32:4743–4754.

Untiet V, Nedergaard M, Verkhratsky A (2024) Astrocyte chloride, excitatory-inhibitory balance and epilepsy. Neural Regen Res 19:1887.

Untiet V, Kovermann P, Gerkau NJ, Gensch T, Rose CR, Fahlke C (2017) Glutamate transporter-associated anion channels adjust intracellular chloride concentrations during glial maturation. Glia 65:388–400.

Verkman AS, Sellers MC, Chao AC, Leung T, Ketcham R (1989) Synthesis and characterization of improved chloride-sensitive fluorescent indicators for biological applications. Anal Biochem 178:355–361.

Yu EP, Dengler CG, Frausto SF, Putt ME, Yue C, Takano H, Coulter DA (2013) Protracted postnatal development of sparse, specific dentate granule cell activation in the mouse hippocampus. J Neurosci 33:2947–2960.

Zhang LL, Pathak HR, Coulter DA, Freed MA, Vardi N (2006) Shift of intracellular chloride concentration in ganglion and amacrine cells of developing mouse retina. J Neurophysiol 95:2404–2416.

